# Microbiota responses to different prebiotics are conserved within individuals and associated with habitual fiber intake

**DOI:** 10.1101/2021.06.26.450023

**Authors:** Zachary C. Holmes, Max M. Villa, Heather K. Durand, Sharon Jiang, Eric P. Dallow, Brianna L. Petrone, Justin D. Silverman, Pao-Hwa Lin, Lawrence A. David

**Affiliations:** Department of Molecular Genetics and Microbiology, Duke University, Durham, NC; Center for Genomic and Computational Biology, Duke University, Durham, NC; Medical Scientist Training Program, Duke University, Durham, NC; College of Information Science and Technology, Penn State University, University Park, PA; Department of Medicine, Penn State University, Hershey, PA; Institute for Computational and Data Science, Penn State University, University Park, PA; Duke Molecular Physiology Institute, Duke University, Durham, NC; Department of Medicine, Duke University Medical Center, Durham, NC; Program in Computational Biology and Bioinformatics, Duke University, Durham, NC

**Keywords:** Microbiome, prebiotic, short-chain fatty acids, diet, personalized nutrition, fiber

## Abstract

**Background:** Short-chain fatty acids (SCFAs) derived from gut bacteria are associated with protective roles in diseases ranging from obesity to colorectal cancers. Intake of microbially accessible dietary fibers (prebiotics) lead to varying effects on SCFA production in human studies, and gut microbial responses to nutritional interventions vary by individual. It is therefore possible that prebiotic therapies will require customizing to individuals.

**Results:** Here, we explored prebiotic personalization by conducting a three-way crossover study of three prebiotic treatments in healthy adults. We found that within individuals, metabolic responses were correlated across the three prebiotics. Individual identity, rather than prebiotic choice, was also the major determinant of SCFA response. Across individuals, prebiotic response was inversely related to basal fecal SCFA concentration, which, in turn, was associated with habitual fiber intake. Experimental measures of gut microbial SCFA production for each participant also negatively correlated with fiber consumption, supporting a model in which individuals’ gut microbiota are limited in their overall capacity to produce fecal SCFAs from fiber.

**Conclusions:** Our findings support developing personalized prebiotic regimens that focus on selecting individuals who stand to benefit, and that such individuals are likely to be deficient in fiber intake.

## Background

A key beneficial role of the human gut microbiota is producing the short-chain fatty acids (SCFAs) acetate, propionate, and butyrate, which are metabolic end products of anaerobic fermentation (1, 2). These SCFA, most notably among them, butyrate, have been shown to promote gastrointestinal (GI) barrier integrity, lower inflammation, increase resistance to pathogen invasion, and contribute positively to enteroendocrine signaling and energy homeostasis (3–9). Augmenting these functions may explain the protective role of SCFAs in diseases ranging from obesity to inflammatory bowel diseases (3, 10–20).

The substrates for SCFA production in the human gut are predominately fibers, which are dietary carbohydrates inaccessible to human enzymes and transporters (21). On average, American adults consume only 21 to 38% of the fiber that is recommended by the USDA (22). This lack of fiber is implicated in the etiology of Western diseases (15, 23, 24), which may in part be tied to the effects of low fiber consumption on microbial SCFA production and SCFA signaling (15, 25–29). To address fiber deficiencies, dietary supplements known as prebiotics have been formulated using specific dietary fibers that are microbially fermentable but host-inaccessible. Prebiotics can increase production of SCFAs by the gut microbiota (30–33), and are a promising microbiota-targeting approach for promoting health (34).

A key challenge of using prebiotics to promote SCFA production is heterogeneity in treatment response. Many distinct prebiotic formulations exist (35, 36), and different prebiotics have been associated with diverging patterns of SCFA production (30, 32). Prebiotic trials have also often revealed an increase in SCFA at the population level following treatment, but inconsistent results at the level of individual (30–33). Still, prebiotic trials tend not to examine how responses to different prebiotics vary within individuals, and have to date often profiled single prebiotics (31, 33, 37, 38), or tested different prebiotics in separate cohorts (30, 32, 39). More rare crossover studies have taken place, but have either not investigated individual variation in prebiotic response (40), or solely utilized *in vitro* assays of prebiotic response (41, 42). If consistent evidence does emerge that responses to different prebiotics vary within people, then effective prebiotic therapies will face the challenging task of personalizing prebiotic choices for each recipient. Alternatively, if responses to different prebiotics are conserved within individuals, effective prebiotic therapy could primarily focus on selecting individuals for treatment. Evidence supporting this latter scenario include studies showing that individual identity is a stronger predictor of SCFA concentrations than dietary intervention (43), and that individual characteristics like habitual food choice can predict changes to microbiota community composition following prebiotic supplementation (38).

To directly investigate the interaction of prebiotic and individual variation in prebiotic response, we report here the results of a randomized, three-way, crossover trial where a single set of individuals was treated with multiple prebiotics. The design was uniform in period (each intervention occurring the same number of times in an intervention period) and in series (each intervention occurring the same number of times in a intervention series), and fully balanced with respect to carryover effects (each intervention is preceded by each other intervention an equal number of times across the study). Our primary lines of inquiry were to assess the relative contribution of individual and prebiotic differences to SCFA responses to supplementation, with an emphasis on butyrate production. Secondary analyses included determining what host and microbiological factors shaped variation in prebiotic response.

## Methods

### Participant recruitment

This study was conducted at Duke University in Durham, NC, USA, and was registered with ClinicalTrials.gov (Identifier: <u>NCT03595306</u>). All recruitment and study procedures involving human subjects were approved by the Duke University Institutional Review Board (Pro00087214). Study participants were recruited by flyers, email list-servs, and word-of-mouth at Duke University Campus. Individuals with a self-reported history of irritable bowel syndrome, inflammatory bowel disease, type-2 diabetes, kidney disease, intestinal obstruction, or colorectal cancer were excluded. Participants were also excluded if they had taken oral antibiotics within the past six months, had any known food allergies, reported dietary intolerances of any kind, or were pregnant or breastfeeding. If individuals identified as female, their urine was subjected to a pregnancy test (Quidel QuickView One-Step hCG Urine Test) to minimize the risk that prebiotic treatment would impact newborn or infant health. Participants who tested positive were deemed ineligible for participation in the study. Inclusion criteria were that participants were between the ages of 18 and 70, and able to provide stool samples at no risk to themselves. Fiber consumption, prebiotic/probiotic usage, and dietary habits were not used as inclusion/exclusion criteria, but participants were asked to report via surveys as part of the study requirements. During this period, 81 individuals set up a phone interview to determine eligibility and interest to participate in the study. Of these, 41 completed electronic informed consent to the study and were subsequently enrolled. Ultimately, eight subjects were lost to follow-up, two withdrew, and three were omitted from some or all analyses due to inability to collect sufficient usable sample, resulting in 28 subjects included in all final data analyses.

### Study design

The three-period crossover longitudinal study was conducted from May of 2018 to March of 2019. The crossover design allowed investigation of three prebiotic interventions while also allowing participants to serve as their own controls (Figure 1). There were six arms of the study, to cover all possible orders of prebiotic consumption and to eliminate the need to account for uneven carry-over effects. The study consisted of an initial study visit, followed by a 6-week study period with alternating weeks of baseline (no prebiotic consumption) and intervention (prebiotic consumption). Participants were added sequentially to an arm group as they enrolled; meaning the first participant was enrolled in Arm one, the second to Arm two and so on. Participants were blinded to the prebiotics that they were consuming until the end of the study. The prebiotics were taken twice a day, Monday to Friday during the intervention weeks. Monday was a half-dose, to help allow participants to adjust to the increase in fiber. Following consent, participants had an initial study visit, to have their baseline physiology assessed by height, weight, and blood pressure measurement, as well as provide a urine sample and receive the study kit and materials. Participants were asked to self-collect stool at home three times a week each week during the study period. Participants also completed four diet surveys throughout the study; one instance of the Diet History Questionnaire Version 3 (DHQ3) (44), covering the one month period prior to the study, and three instances of the Automated Self-Administered 24-hour (ASA24) Dietary Assessment Tool, version 2017, developed by the National Cancer Institute, Bethesda, MD (45). Two ASA24’s were completed during the week, and one on a weekend. The DHQ3 was completed prior to the 6-week study period. We found that correlations of total calories, protein, fat, carbohydrate, and dietary fiber between the two tools were robust (Calories; p = 0.032, r = 0.42: Protein; p = 0.021, r = 0.45: Fat; p = 0.036, r = 0.41: Carbohydrates; **p = 0.069**, r = 0.38: Dietary fiber; p = 0.001, r = 0.59: Pearson correlations). Additionally, participants completed online-surveys about their experiences with the prebiotics following the intervention weeks.

**Figure 1:**
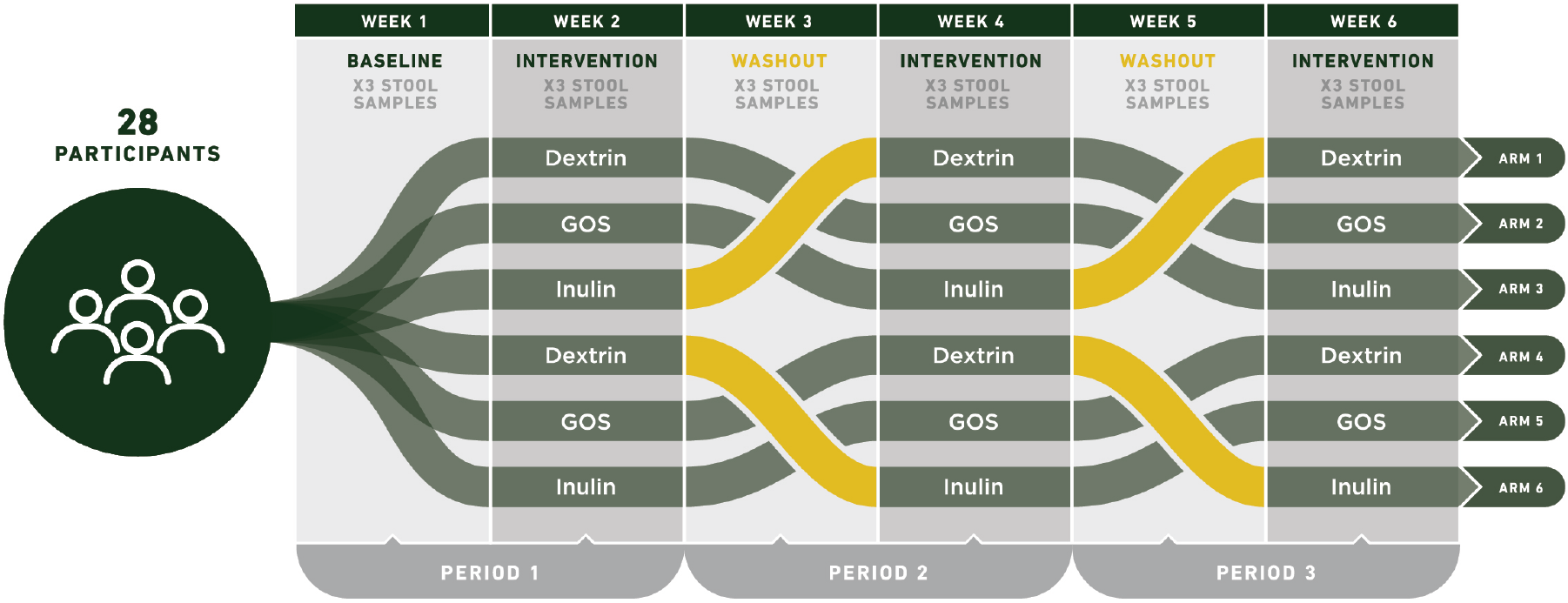
Prebiotic trial design. A three-period crossover design was standardized in series and period, with each prebiotic occurring the same number of times within each arm (once) and study period (twice). The design was also balanced with respect to first-order carry-over effects, meaning prebiotics were administered in all possible orders. Each of the three periods consisted of a prebiotic-free week followed by a prebiotic-supplemented week. Six study arms encompassed all possible orders of prebiotic supplementation, and subjects were assigned to arms sequentially by time of enrollment, resulting in approximately equal numbers of individuals in each arm. A total of 41 participants were enrolled and 28 were considered to have completed the study.

### Prebiotic supplementation

Three prebiotics were tested in this study: inulin (manufactured by Now Foods, Bloomingdale, IL), wheat dextrin (Benefiber, manufactured by GSK Consumer Healthcare, Warren, NJ), and galactooligosaccharides (GOS) (Bimuno, manufactured by Clasado Limited, Shinfield, UK). In a clean non-laboratory space, prebiotics were weighed into individual doses and packaged into pre-labeled food-safe sealable plastic bags. Each participant received all three prebiotics throughout the study; inulin at 9g / day, wheat dextrin at 9g / day, and galactooligosaccharides (GOS) at 3.6g / day. These doses correspond to the recommended daily intake on the product label. In order to further minimize the possibility of discomfort associated with consuming prebiotics, the first day of dosing for each prebiotic was a half dose to allow acclimation (4.5g for inulin and dextrin, 1.8g for GOS). Participants consumed one of the three prebiotics at a time, twice a day, for a duration of five days, followed by a week of washout. The order was determined by their assigned study arm. Participants were instructed to consume the prebiotics by mixing the powders in any kind of beverage (such as water) or food (such as applesauce), so long as it was cold or room temperature.

### Compliance and side effects

Surveys suggested participants complied with the design of the prebiotic intervention. After each prebiotic intervention week (weeks 2, 4, and 6), participants were sent a survey that asked about missed doses. 26 of 28 participants responded to the surveys, with an average of 2.77 out of 3 surveys being returned. Of the 72 person-weeks surveyed, 18 included missed doses. Of 720 doses surveyed, 26 were missed, indicating a study average missed dose rate of 3.6%. No respondent missed more than 14% of doses over the course of the study, and none missed more than 30% of doses during any intervention week.

Side-effects reporting also provided evidence that participants consumed the prebiotics. After each prebiotic intervention week, participants were sent a survey that asked about side effects. When asked to identify what extent they agree with the statement “The prebiotic caused my gastrointestinal discomfort, 7 out of 23 respondents indicated that they ‘agree’ with this statement (as opposed to ‘neither agree nor disagree’, ‘disagree’, and ‘strongly disagree’) during the inulin supplementation week. For GOS and dextrin, 3 of 25 and 4 of 24 indicated agreement. When asked whether the prebiotic affected the specific symptoms of abdominal pain, bloating, flatulence, nausea, reflux/heartburn, and borborygmi, fewer than 20% of participants reported ‘increased’ or ‘greatly increased’ symptoms, except in the case of flatulence (all prebiotics) and abdominal pain (inulin). 12 of 23 respondents indicated increased or greatly increased flatulence on inulin, 6 of 25 on GOS, and 8 of 24 on dextrin. Six of 23 respondents indicated increased abdominal pain on inulin, compared of none on GOS and 2 on dextrin. There were no differences in self-reported weekly average Bristol stool hardness between baseline and any of the three prebiotic interventions (p = 0.61, 0.20, and 0.38 for inulin, GOS, and dextrin, respectively; paired t tests).

Consistent with the notion that prebiotic impacts are stronger in individuals who tend to consume less fiber, we found evidence of a higher likelihood for side effects in those individuals who habitually consumed less fiber. Participants who indicated that GOS or Dextrin increased (‘greatly increased’ or ‘increased’) flatulence had significantly lower habitual fiber consumption than did those participants who indicated that GOS or Dextrin decreased or did not affect flatulence (‘did not affect’ or ‘reduced’) (p = 0.006 and p = 0.045 for GOS and Dextrin, respectively; t-test). This affect was not present for inulin (p = 0.45). Similarly, Participants who indicated that GOS or Dextrin increased (‘greatly increased’ or ‘increased’) borborygmi had significantly or nearly significantly lower habitual fiber consumption than did those participants who indicated that GOS or Dextrin decreased or did not affect borborygmi (‘did not affect’ or ‘reduced’) (p = 0.065 and p = 0.019 for GOS and Dextrin, respectively; t-test). This affect was not present for inulin (p = 0.64). Further, fiber consumption other than the provided supplement did not change over the course of the study (p = 0.15, ANOVA).

### Stool collection

At the pre-study visit, participants were provided with a standard urine specimen collection cup and shown to a bathroom to collect the sample. Urine samples were also sent to LabCorp in a BD Vacutainer UA tube to measure metabolic variables, including routine urinalysis, sodium, potassium, urea nitrogen, and creatinine. Consented participants were asked to self-collect stool three times a week at home using sampling kits provided to them. Participants were asked to immediately store samples in their personal freezers (−20C). Participants were asked to bring stored samples to the David Lab freezers on a weekly basis. Stool samples were transferred and stored at −80C until thawed for DNA extraction and other sample processing.

### *In vitro* fermentation

*In vitro* fermentation of stool microbiota was conducted following protocols previously published (41). This method has been shown to be robust to the effects to the effects of freezing stool, with total SCFA production remaining correlated between fresh stool and stool frozen and thawed up to two times (41), and butyrate production remaining correlated between fresh stool and stool frozen once (p = 0.0004, ρ = 0.49, Spearman correlation), as were samples in this study. In brief, a 10% w/v fecal slurry was made using thawed week-1 stool and anaerobic 1X PBS (pH 7.0 ± 0.1). This slurry was filtered through 0.33mm pore size mesh, then mixed 1:1 with a 1% w/v solution of prebiotic and incubated in duplicate in a 24-well cell culture plate for 24hr. We expect these *in vitro* fermentations to represent the non-adherent microbial fraction of stool, as well as bacteria adhered to smaller particles that pass through a 0.33mm filter. Three prebiotic treatments were used, corresponding to the three prebiotics used in the *in vivo* portion of this study: inulin (Now Foods Inulin Powder, part #2944), galactooligosaccharides (GOS; Bimuno Powder), and wheat dextrin (Benefiber Original). The resulting fermentation conditions where therefore 5% fecal slurry with 0.5% prebiotic (w/v) and occurred at 37°C for 24 hours. Following fermentation, a 1mL aliquot was taken from each well for SCFA quantification.

### Quantification of SCFA

Quantification of SCFA by gas chromatograph flame ionization detection (GCFID) was performed following protocols previously published (41). In brief, a 1mL aliquot of either 10% fecal slurry in PBS or a 10% solution of the fermentation vessel contents was obtained. To this, 50 μL of 6N HCl was added to acidify the solution to a pH below 3. The mixture was vortexed, centrifuged at 14,000rcf for five minutes at 4°C to remove particles. Avoiding the pellet, 750 μL of this supernatant was passed through a 0.22μm spin column filter. The resulting filtrate was then transferred to a glass autosampler vial (VWR part #66009-882).

Filtrates were analyzed on an Agilent 7890b gas chromatograph (GC) equipped with a flame-ionization detector (FID) and an Agilent HP-FFAP free fatty-acid column (25m x .2mm id x .3μm film). Acetate, propionate, isobutyrate, butyrate, isovalerate, and valerate were identified and quantified in each sample by comparing to an 8-point standard curve that encompassed the sample concentration range. Standards contained 0.1mM, 0.2mM, 0.5mM, 1mM, 2mM, 4mM, 8mM, and 16mM concentrations of each SCFA. Once measured, the SCFA concentrations were multiplied by a scaling factor of 10 to return them to their undiluted value. We report SCFA concentrations as mM, however, based on our methodology using a standardized stool mass / diluent ratio, these mM measurements are quantitatively equivalent to mmol/kg (*e.g.* 5mM is equal to 5mmol/kg).

### DNA extraction, PCR amplification, and sequencing

We performed 16S rRNA gene amplicon sequencing on human stool samples to determine microbiota community composition following protocols previously published (41). DNA was extracted from frozen fecal samples with the Qiagen DNeasy PowerSoil DNA extraction kit (ID 12888-100). Amplicon sequencing was performed using custom barcoded primers targeting the V4 region of the 16S rRNA gene (46), using published protocols (46–48). The sequencing library was diluted to a 5pM concentration and sequenced using an Illumina MiniSeq and a MiniSeq Mid Output Kit (FC420-1004) with paired-end 150bp reads.

### Identifying sequence variant and taxonomy assignment

We used an analysis pipeline with DADA2 (49) to identify and quantify sequence variants, as previously published (41, 50). To prepare data for denoising with DADA2, 16S rRNA gene primer sequences were trimmed from paired sequencing reads using Trimmomatic v0.36 without quality filtering (51). Barcodes corresponding to reads that were dropped during trimming were removed using a custom python script. Reads were demultiplexed without quality filtering using python scripts provided with Qiime v1.9 (52). Bases between positions 10 and 150 were retained for the forward reads and between positions 0 and 140 were retained for the reverse reads. This trimming, as well as minimal quality filtering of the demultiplexed reads was performed using the function fastqPairedFilter provided with the DADA2 R package (v1.8.0). Sequence variants were inferred by DADA2 independently for the forward and reverse reads of each of the two sequencing runs using error profiles learned from all 20 samples. Forward and reverse reads were merged. Bimeras were removed using the function removeBimeraDenovo with default settings. Taxonomy was assigned using the function assignTaxonomy from DADA2, trained using version 123 of the Silva database.

### Statistics and modelling

All statistical analyses and modelling for this study were conducted in R (R version 3.4). T-tests, Wilcoxon Rank-Sum tests, Spearman correlations, Pearson correlations, and ANOVA tests were conducted with base R. PERMANOVAs were carried out with the ‘adonis2’ function from the R package ‘vegan’. Differential abundance tests were carried out using the R package ‘ALDEx2’. Linear models, and generalized linear models were implemented with the ‘lm’ and ‘glm’ functions from base R. Generalized linear mixed models (GLMM) were implemented with the ‘glmer’ function from R package ‘lme4’. P values for effects in linear models and generalized linear models were calculated using the ‘summary’ function in base R on fitted ‘lm’ and ‘glm’ objects and are based on F-tests. P values for effects in GLMMs were calculated using the ‘summary’ function (base R) on fitted ‘glmer’ objects, and are based on Wald Z-tests. All R code used to analyze the data presented in this manuscript is freely available on a publicly accessible repository (https://doi.org/10.6084/m9.figshare.c.5405505).

## Results

### Study Participants

We enrolled participants into a six-week, three-period prebiotic intervention study (Figure 1). We used a crossover design to compare multiple prebiotics across a subject pool. We selected inulin, wheat dextrin, and galactooligosaccharides (GOS) as our prebiotics, as these are commonly consumed (53), commercially available dietary supplements, and are monomerically distinct. Our crossover was standardized in series and period, with each prebiotic occurring the same number of times within each series (once) and study period (twice). The design was also balanced with respect to first-order carry-over effects, meaning we shuffled the order in which prebiotics were administered so as to minimize the interactive effects between any two pairs of consecutive prebiotics. Each study period consisted of an intervention-free baseline week and a prebiotic intervention week. Stool samples were collected on days three, four, and five of each of the six study weeks (baselines and intervention). Of the 41 participants who initially enrolled, two withdrew, three were unable to collect at least one usable sample (i.e., insufficient volume or inadequate labeling) per week, and eight were lost to follow-up. We focused our subsequent analyses on the remaining 28 participants who completed the study. These participants in total collected 413 stool samples (mean=2.46 stool samples per person per week).

### Prebiotic supplementation affected SCFA production at the cohort-level

We first sought to examine differences in microbiome response during each of the different prebiotic treatments. Analyses of gut microbial community composition did not reveal associations with inulin, GOS, or wheat dextrin treatment (p > 0.05, PERMANOVA using unweighted UniFrac distance at species level). In addition, no species were statistically differentially abundant during any prebiotic intervention and its corresponding prebiotic-free baseline (p > 0.05, Benjamini-Hochberg corrected Wilcoxon Rank Sum tests using the R package ALDEx2). While previous studies have observed changes in community composition following prebiotic administration, such effects have been observed in a subset of tested prebiotics (30, 32), and with different dosing schedules (30–32, 38). We also did not observe lasting shifts in community composition over the course of the study at the cohort level, as baseline weeks three and five did not differ from baseline week one by PERMANOVA (p = 0.995 and 0.999, respectively).

Still, in contrast to the lack of a coherent 16S rRNA gene shifts among participants, we found the three prebiotics varied according to resulting fecal SCFA profiles relative to baseline. GOS was associated with decreased total stool SCFA concentrations (Table 1, Figure S1C, effect size = −0.071, p = 0.038, GLMM), an outcome that has been previously observed (54). Differences within individual SCFAs were also apparent when analyzing absolute fecal SCFA concentrations (Table 1). Dextrin increased propionate concentration (effect size = 0.074, p = 0.038, GLMM), and GOS decreased both propionate and butyrate concentrations (effect size = −0.087, p = 0.015; effect size = −0.14, p = 0.002, GLMMs, respectively). Still, changes in total and individual SCFA concentrations may be caused by effects of stool hydration, which can vary up to three-fold and directly impacts measures of SCFA concentration in feces (55, 56). To assess the relative production of each SCFA, we therefore normalized levels of individual SCFAs to account for differences in stool hydration. Our resulting analysis found that inulin supplementation increased proportional butyrate concentrations (butyrate concentration / total SCFA concentration) in participants’ stool (effect size = 0.082, p = 0.007, GLMM; Figure 2, Table 1). These findings are consistent with prior reports that inulin supplementation may increase butyrogenesis (57–59). Our analyses also suggested GOS and wheat dextrin decreased proportional butyrate concentration in participants’ stool (GOS; effect size = −0.074, p = 0.012. Dextrin; effect size = −0.071, p = 0.015, GLMM; Figure 2, Table 1). These findings align with prior trials that found both GOS (54, 60, 61) and dextrin (62, 63) increases acetate production over more reduced SCFAs, including butyrate and propionate, potentially by shunting nutrients away from butyrate production. No prebiotic altered proportional acetate or proportional propionate concentration at the cohort level (p > 0.05, GLMM; Figures S1A and S1B, Table 1).

**Figure 2:**
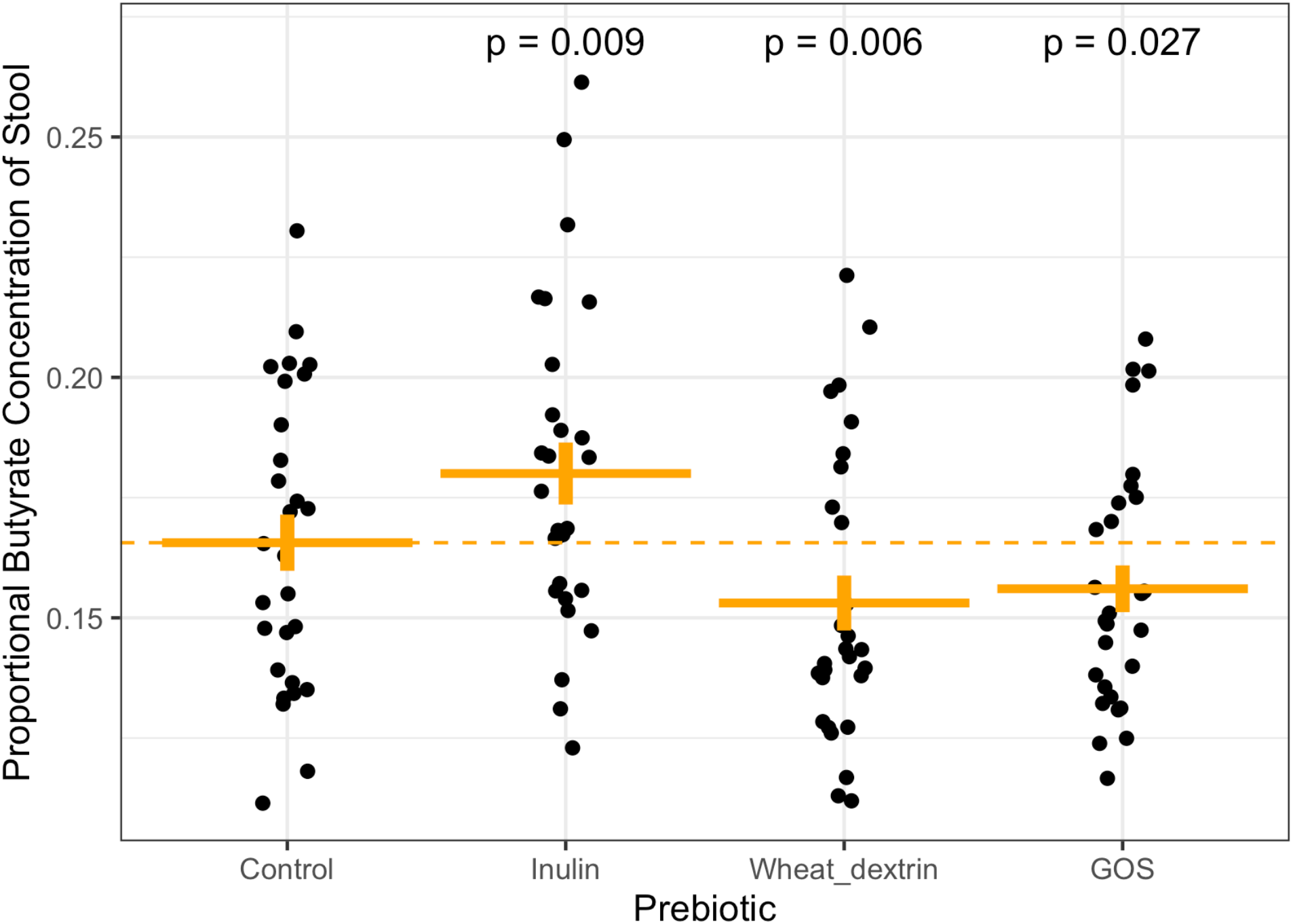
Effects of prebiotic supplementation on proportional butyrate concentration of stool. Plotted are the average proportional butyrate concentrations for each study participant during each prebiotic intervention period (up to three samples per person per week), and the average of all three baseline weeks (up to nine samples per person). Inulin was associated with significantly increased proportional butyrate concentration of stool averaged across individuals, while both dextrin and GOS were associated with decreased proportional butyrate relative to control (dashed line) (generalized linear mixed model). A proportional butyrate concentration value of 1 = 100%.

**Table 1:** Concentrations and proportions of SCFA in stool during prebiotic and control weeks, averaged across individuals. Significance was determined by generalized linear mixed models (see methods). Concentrations of stool expressed as mM are equivalent to mmol/kg (see methods).

|  | Control | Inulin |  | GOS |  | Dextrin |  |
| --- | --- | --- | --- | --- | --- | --- | --- |
|  | Median<br>[min –<br>max] | Median<br>[min –<br>max] | p | Median<br>[min –<br>max] | p | Median<br>[min –<br>max] | p |
| **CONCENTRATION** |  |  |  |  |  |  |  |
| **Acetate (mM)** | 47.4<br>[5.6 –<br>102.1] | 46.9<br>[11.6 –<br>94.1] | 0.52 | 47.2<br>[14.8 –<br>80.0] | 0.18 | 50.6<br>[10.9 –<br>94.7] | 0.39 |
| **Propionate (mM)** | 13.1<br>[3.2 –<br>37.8] | 13.1<br>[5.7 –<br>21.8] | 0.41 | 12.3<br>[2.6 –<br>26.0] | **0.015** | 14.4<br>[5.7 –<br>29.6] | **0.038** |
| **Butyrate (mM)** | 12.1<br>[2.8 –<br>40.3] | 12.4<br>[4.1 –<br>34.5] | 0.26 | 10.2<br>[4.1 –<br>29.5] | **0.002** | 10.9<br>[4.6 –<br>31.3] | 0.29 |
| **Total SCFA (mM)** | 75.1<br>[15.6 –<br>153.7] | 72.7<br>[27.9 –<br>131.1] | 0.67 | 68.6<br>[25.6 –<br>114.4] | **0.038** | 76.4<br>[21.1 –<br>145.4] | 0.51 |
| **PROPORTION** |  |  |  |  |  |  |  |
| **Acetate (%)** | 64.6<br>[41.9 –<br>77.9] | 63.7<br>[22.1 –<br>74.6] | 0.21 | 65.9<br>[45.9 –<br>86.8] | 0.19 | 65.7<br>[44.5 –<br>74.9] | 0.43 |
| **Propionate (%)** | 18.4<br>[9.6 –<br>36.2] | 19.6<br>[8.1 –<br>34.7] | 0.63 | 18.8<br>[4.5 –<br>35.4] | 0.70 | 20.1<br>[10.3 –<br>36.4] | 0.31 |
| **Butyrate (%)** | 16.1<br>[8.0 –<br>32.9] | 18.0<br>[11.2 –<br>55.1] | **0.009** | 15.3<br>[8.6 –<br>28.0] | **0.027** | 14.6<br>[10.4 –<br>30.4] | **0.006** |

### Butyrogenic responses are more strongly predicted by individual identity than by prebiotic

We next investigated how prebiotic responses varied among individuals. We focused our analysis on prebiotic impacts on butyrate production because it was the sole SCFA that was altered at the cohort-level, and it is also the microbially derived SCFA most commonly implicated in enteric health (2, 6–8, 64–67). We also focused on proportional butyrate concentrations to avoid the effects of stool hydration on our analysis and to be consistent with previous individual-level analyses of fecal SCFAs (32, 43). In line with prior studies of prebiotic response, (41) statistical modeling yielded evidence for individual-specific butyrogenic responses to each of the prebiotics (that is, an interaction between individual and prebiotic terms in a GLMM). Nonetheless, further analyses suggested that the strongest determinant of proportional butyrate responses was shaped by individual identity: individual-specific effects (pseudo-R^2^ = 0.39, p < 0.0001; GLM F-test) were much stronger than prebiotic-specific effects (pseudo-R^2^ = 0.05, p = 0.001; GLM F-test). Additionally, individuals who exhibited the largest response in proportional stool butyrate concentration on inulin were more likely to exhibit larger responses on dextrin (ρ = 0.45, p = 0.015, Spearman correlation; Figure 3). We observed a similar, positive, non-significant correlation between dextrin and GOS (ρ = 0.35, p = 0.065). This relationship was not observed for inulin and GOS (ρ = 0.12, p = 0.53). This pattern was apparent in spite of correcting for baseline differences in SCFA production between individuals, which were also consistent over time and between baseline and treatment weeks.

**Figure 3:**
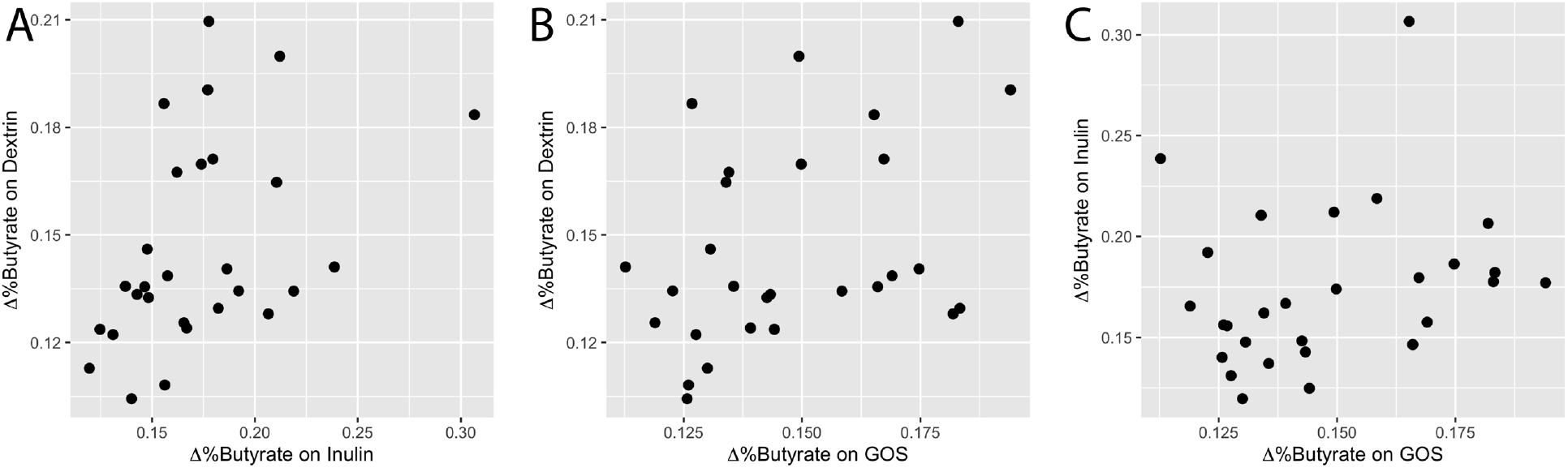
Individual changes in proportional butyrate concentration (Δ%butyrate) compared across multiple prebiotics. Changes in Δ%butyrate were positively correlated when comparing responses to dextrin and inulin (ρ = 0.45, p = 0.015; Spearman correlation), as well as responses to dextrin and GOS (ρ = 0.35, p = 0.065). The relationship between inulin and GOS response trended towards a positive association (ρ = 0.12, p = 0.53). A proportional butyrate concentration value of 1 = 100%.

### Individuals consistently varied according to SCFA metabolic state, microbiota community composition, and SCFA production potential

Why might individual-specific effects be stronger than prebiotic-specific effects in our cohort? We observed that individuals were idiosyncratic in their SCFA profiles over time. Both proportional butyrate concentration and butyrate concentration were correlated within individuals between the first and last baseline weeks (ρ = 0.49, p < 0.007; and ρ = 0.63, p = 0.0003, respectively, Spearman correlations; Figure S2). Variation between individuals’ SCFA profiles could also be linked to inter-individual microbiota variation, a well-known factor in prebiotic response (43). When we divided participants into ‘high’ (> median) and ‘low’ (<= median) fecal butyrate groups, we observed a significant effect of this grouping on microbiota community composition (R^2^ = 0.075, p = 0.0022, PERMANOVA using unweighted UniFrac distance; Figure S3). We identified three specific genera that were differentially abundant between the groups: *Anaerofilum* and *Ruminiclostridium_5* were decreased in the high butyrate group, and *Lachnospira* was increased in the high butyrate group (p = 0.009, .043, and 0.030, respectively; Benjamini-Hochberg corrected Wilcoxon Rank Sum tests; Figure S3). We additionally observed a relationship between baseline microbiota community structure and average response to all three prebiotics (Δ%butyrate). After dividing participants into ‘responders’ and ‘non-responders’, based on whether average Δ%butyrate was positive or negative, we found a significant effect of this grouping on microbiota community composition (R^2^ = 0.059, p = 0.028, PERMANOVA using unweighted UniFrac distance; Figure S4). Thus, individuals’ starting microbiome composition was associated with prebiotic treatment outcome.

### Baseline intake patterns of dietary fiber negatively correlated with response to prebiotic intervention

Next, we sought to understand what host factors might influence individuals’ microbiomes and ultimately shape their butyrogenic responses across the tested prebiotics. We focused on fiber intake, which has been shown to shape microbiota community composition and metabolic output (68, 69). We measured fiber consumption using a dietary questionnaire spanning the month preceding the study (44), and we compared fiber intakes to each individual’s average response to the three prebiotic interventions (average Δ%butyrate). Despite prior studies suggesting that plant or fiber intake would augment gut microbial metabolic capacity (38, 69–71), our comparison revealed that dietary fiber consumption was negatively correlated with average Δ%butyrate (ρ = −0.40, p = 0.046; Spearman correlation; Figure 4A). Thus, higher habitual fiber intake was associated with a lower average Δ%butyrate during the prebiotic intervention. To corroborate these findings using an objective biomarker, we also compared average Δ%butyrate to baseline spot urinary potassium, a known marker for fruit and vegetable intake (72), as well as for general healthy eating (73, 74). We validated that baseline spot urinary potassium was correlated with dietary fiber intake in our study (ρ = 0.47, p = 0.031; Spearman correlation; Figure S5A). While we found no significant correlation between baseline spot urinary potassium and average Δ%butyrate (ρ = −0.38, p = 0.071; Spearman correlation; Figure S5B), the magnitude and direction of effect suggests a statistical relationship may have been apparent with a larger sample size.

**Figure 4:**
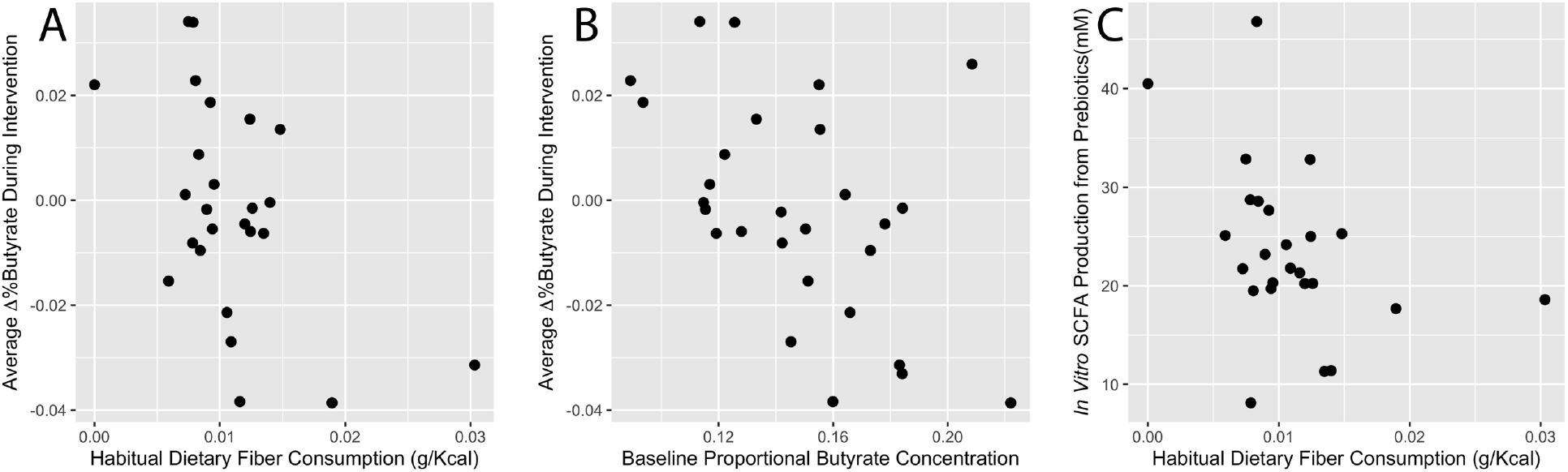
Relationships between habitual diet, baseline proportional butyrate concentration, and metrics of SCFA production. (A) Average change in proportional butyrate concentration of stool during intervention (Δ%butyrate) was negatively correlated with habitual dietary fiber consumption (ρ = −0.40, p = 0.046; Spearman correlation). (B) Average Δ%butyrate was also negatively correlated with baseline proportional butyrate concentration (ρ = −0.49, p = 0.008; Spearman correlation). (C) This relationship between habitual fiber consumption and SCFA production from prebiotics was also observed *in vitro* (ρ = −0.46, p = 0.021; Spearman correlation).

### Observational evidence that baseline SCFA metabolic state and dietary history limits prebiotic response

To understand why habitual fiber intake would be negatively associated with prebiotic responses, we considered recent ecological studies suggesting that nutrient limitation is a key factor shaping the metabolism of mammalian gut microbiota (75, 76). Even if excess fiber is available to polysaccharide utilizing microbes, fixed caps in the availability of other nutrients involved in carbohydrate fermentation could ultimately affect the relative production of different SCFA species. Such relative changes would be caused by differences in the oxidation state of SCFAs. Acetate is the most oxidized SCFA and tends to be produced first and in higher quantities during fermentation (76–78). By contrast, butyrate is the most reduced (76, 78) and the least energetically favorable (76) primary SCFA for the microbe to produce. Thus, changes to overall SCFA production should be associated with changes in proportional butyrate (76–78); and, by extension, individuals who naturally consume high levels of fiber and already activate SCFA production pathways would exhibit reduced capacity to increase proportional butyrate levels following prebiotic treatment.

We identified three findings that were consistent with a threshold model of metabolic capacity limitation. First, we observed that individuals with higher starting proportional concentrations of butyrate exhibited the weakest butyrogenic prebiotic responses. (p = 0.008, ρ = −0.49, Spearman correlation; Figure 4B). In our study, no participant ultimately exhibited butyrate concentrations in excess of 29mM following prebiotic treatment (though we note that higher upper limits (e.g. greater than 40mM) have been previously observed (30)).

A second finding linking background nutrient intake and metabolic limitation involved baseline proportional butyrate concentrations and habitual diet. Dietary fiber consumption was positively correlated with proportional butyrate concentration at baseline week 1, prior to any prebiotic intervention (1g fiber/2000 kilocalorie diet led to an increase in proportional butyrate concentration of 0.010, p = 0.012, beta regression; Figure S6) as has been previously suggested (33, 37, 79). Furthermore, using an ecological technique for identifying relationships between multivariate datasets known as co-inertia analysis (80), we found a significant interaction between baseline proportional SCFA concentrations and food choices(p = 0.031, coinertia analysis). This analysis involved comparing patterns in the abundance of short chain acids and food intake across matched samples (Figure 5A). Coinertia analysis associated the intake of several fiber-rich food groups with baseline proportional butyrate concentrations including legumes, berry fruits, and dark-green vegetables (vectors pointing towards the top-left in Figures 5B, C).

**Figure 5:**
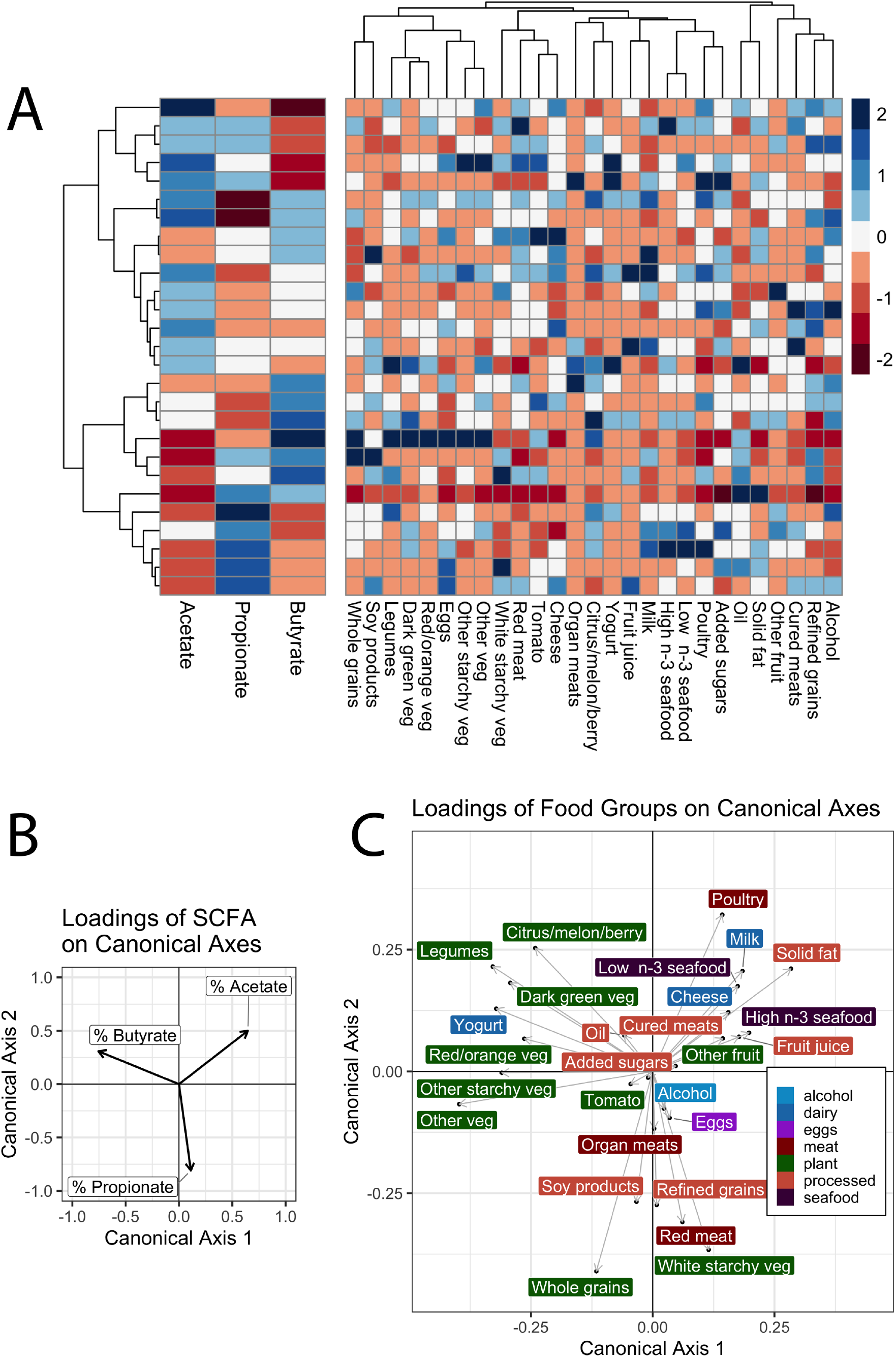
Relationship between baseline SCFA profiles and habitual intake of food groups. (**A**) Baseline SCFA profiles and average food intakes among participants. Participants (rows) clustered based on proportional SCFA profiles. Cells values represent z-scores within each column. Loadings shown of proportional SCFA concentrations (**B**) and consumed food groups (**C**) along canonical axes one and two resulting from co-inertia analysis of SCFA and diet data (p = 0.031). Each axis is represented identically in each plot, so that eigenvectors (direction of arrows) correspond between the SCFA and food group plots and the magnitude of eigenvalues (length of arrows) is related to the strength of that component in the axis loading.

### Culture-based experimental evidence links fiber intake to limits on microbiome metabolic response

A third line of experimental evidence arose linking habitual nutrient intake and microbial SCFA production capacity. We hypothesized that if higher baseline fiber consumption reduced the capacity of gut microbiota to increase proportional butyrate production from prebiotics, the metabolic activity of fecal microbiota would exhibit a negative relationship with fiber intake. To test this hypothesis, we applied a recently developed an *ex vivo* fermentation model (41) to baseline stool samples collected from each study participant. This model experimentally tests the metabolic effects of supplementing a microbial community using a given prebiotic as the sole additional carbon source, independent of potentially confounding host factors like SCFA absorption, and has been shown to be robust to the effects of freezing stool (41). We note that while our *in vivo* measurements rely on proportional SCFA concentrations due to the potentially confounding effects of factors like stool hydration, these *in vitro* measures allow us to directly measure absolute SCFA levels as an indicator of metabolic activity and prebiotic utilization. In support of the model’s capability to produce SCFAs following prebiotic administration, we observed an increase in overall SCFA production relative to control (p < 0.0001, t-test; average of 4.41-fold increase across participants, calculated using the average total SCFA concentration of three separate prebiotic treatments) when the *in vitro* assays were supplemented with prebiotics as the sole carbon source. When we compared SCFA production to fiber intake, we observed as expected a negative correlation between fiber consumption and SCFA production (p = 0.021, ρ = −0.46, Spearman correlation using average of three prebiotic treatments; Figure 4C). We also observed a negative correlation between fiber consumption and SCFA production in prebiotic-free control treatments (p = 0.019, ρ = −0.47, Spearman correlation; Figure S7). Thus, individuals who tended to eat more fiber in our cohort also harbored gut microbiota that produced less SCFA when exposed to prebiotics. Still, we note that a relationship was not observed when *in vitro* %butyrate was compared to baseline fiber consumption (p = 0.37, Spearman correlation). This negative result may reflect the inability of our *in vitro* assays to capture metabolic tradeoffs affecting the relative production of different SCFAs following prebiotic challenge *in vivo*.

### Consistent growth of specific microbial genera supports individual-specific prebiotic response

Next, we used our *ex vivo* fermentations to associate specific gut bacterial taxa with prebiotic response. Following 24hr fermentation, microbial community composition was assayed from each prebiotic-supplemented culture and compared to no-carbon-added control cultures, for each of the 28 participants, using ALDEx2. This analysis revealed multiple genera that were over or under-represented in each prebiotic treatment (Figure 6).

**Figure 6:**
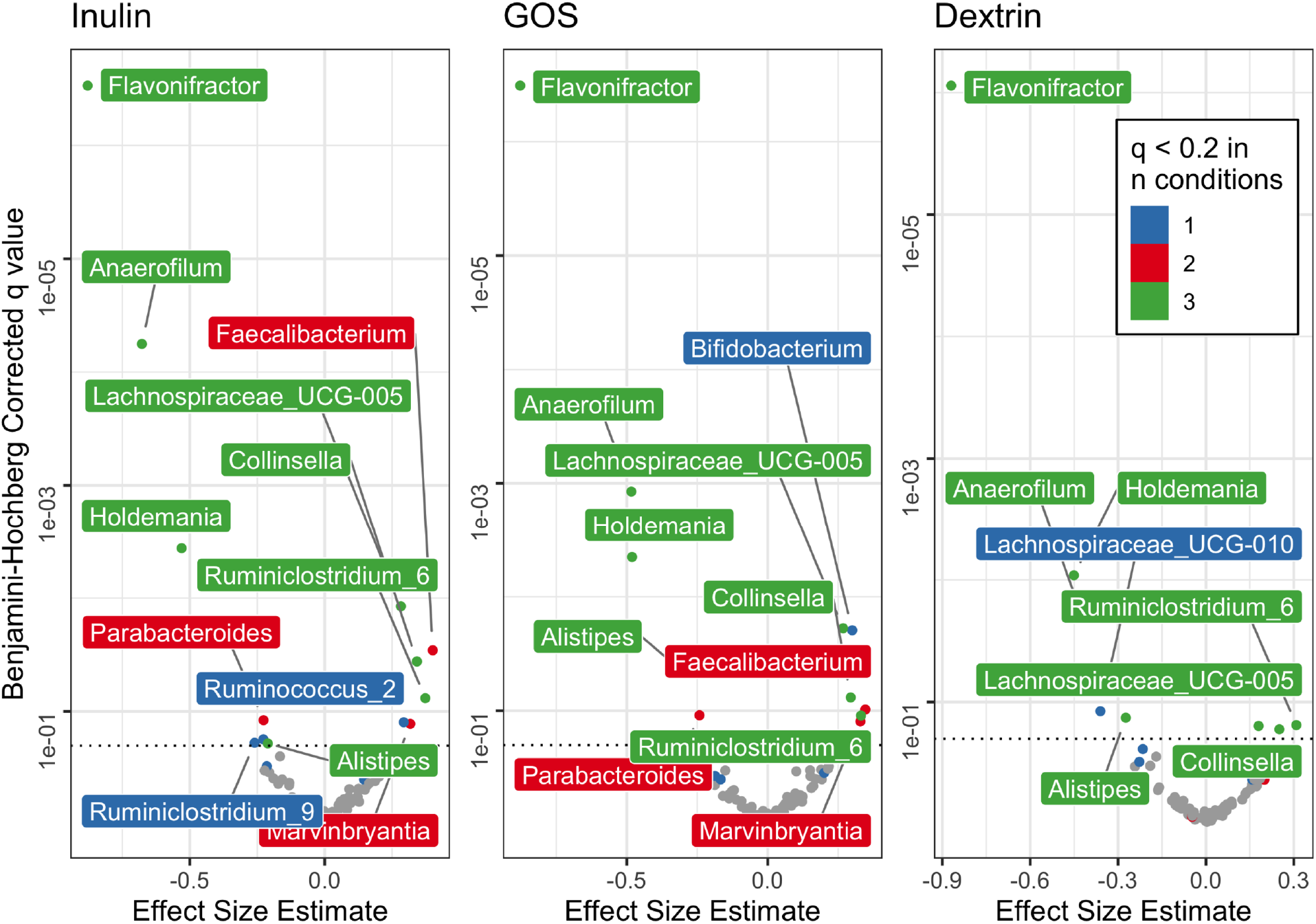
Gut bacterial taxa whose abundance changes during prebiotic-supplemented *ex vivo* fermentation. Bacterial abundances were compared to prebiotic-free control fermentation. Colors represent the number of prebiotic conditions in which a genus was differentially abundant (Benjamini-Hochberg corrected q value < 0.2, visualized with horizontal dashed line). Effect size estimate as reported by ALDEx2 is the median difference between groups (prebiotic and control) divided by the maximal difference within either group, in clr-transformed coordinates.

Additionally, most genera that were differentially abundant in one prebiotic treatment exhibited similar differential abundance in the other prebiotic treatments. This result is consistent with prior studies finding that fiber-degrading bacteria capable of degrading a breadth of dietary carbohydrates (“generalists”) are abundant in the gut (81). Indeed, all genera exhibiting conserved higher abundance in two or three prebiotic treatments compared to control (*Collinsella*, *Faecalibacterium*, *Lachnospiraceae_UCG-005*, *Marvinbryantia*, and *Ruminiclostridium*) are known degraders of common dietary fibers with broad specificity (82–84). By contrast, of the genera exhibiting conserved lower abundance in two or three prebiotic treatments compared to control (*Alistipes*, *Anaerofilum*, *Flavonifractor*, *Holdemania*, and *Parabacteroides*), we are not aware of reports linking them to degradation of inulin, dextrin, or GOS, and only *Parabacteroides* has been shown to degrade any dietary fiber (82, 84). We do note though that we did not observe the aforementioned genera to be associated with prebiotic responses in our *in vivo* data. Such findings suggest that other behavioral or lifestyle factors not captured by *in vitro* assay contribute to variation in microbiome response to prebiotics in humans.

Overall, the consistent response of select bacterial taxa to prebiotic addition *in vitro* further supports the conclusion that responses across a range of prebiotics will be linked to individual-specific factors like gut microbial community composition.

## Discussion

In this study, we administered the prebiotics inulin, GOS, and dextrin to a set of individuals using a fully balanced crossover design. We found that these prebiotics varied in their impact on fecal SCFA concentrations when data was agglomerated at the population scale, with inulin increasing proportional butyrate and both GOS and wheat dextrin decreasing proportional butyrate. Yet, our analysis also suggested that individual identity exerted a much stronger effect on prebiotic response, and we observed that responses to each prebiotic were correlated within individuals. Individual response variation was associated with differences in baseline microbiological and dietary factors. In particular, we found that proportional butyrogenic response to prebiotics was anti-correlated with habitual fiber intake. We confirmed the involvement of the microbiota (as opposed to the host) in this relationship using *in vitro* fermentation experiments. This negative relationship may be explained by limitations in the butyrate production capacity of human gut microbiota.

From a microbial ecological perspective, our findings support a model where microbial metabolic repertoires within individuals’ microbiota have the capacity to utilize a diverse array of chemically distinct fiber types. Evidence in favor of this model include *in vitro* prebiotic screens suggesting that individuals uniformly harbor bacteria capable of degrading a wide variety of carbohydrate species (81). Such metabolic plasticity may benefit gut microbiota by enhancing community stability and invasion resistance (85–87), and could be enabled by substrate-switching at the level of strains (87, 88) or cross-feeding interactions (89). Still, differences in the levels of strains underlying these metabolic interactions may vary between individuals (81, 90), which could in turn lead to inter-individual variation in the capacity of individuals’ microbiota to ferment prebiotics.

The nutritional implications of our work were, in our view, more surprising. A positive relationship between dietary fiber consumption and microbiota community structure (91), and metabolic function (69, 92–94) is well established, with higher fiber intake linked to greater SCFA and, specifically, butyrate production. Thus, we expected that individuals habituated to a high-fiber diet would have a greater capacity to utilize incoming novel fiber towards increasing proportional butyrate production, not a lower capacity as we observed. Still, prior studies reporting links between diet patterns and microbiota response have specifically associated changes to microbial community composition with baseline fiber intake (38). Associations between fiber consumption and SCFA production, by contrast, have either not been tested for in prebiotic trials (95, 96) or not revealed a positive relationship (38). Additionally, while our analyses confirmed an association between baseline proportional butyrate levels and intake of plant foods groups linked to dietary fiber (e.g. legumes and dark green vegetables, Figure 5), it also linked food groups typically associated with a Mediterranean diets (e.g. seafood, dairy, fruit, and poultry (97)) with higher baseline levels of proportional acetate (Figure 5).

In light of the surprising nature of our findings, it is necessary to fully explore other potential drivers of this negative relationship between baseline fiber consumption and prebiotic response. One major concern is the potential for biased study compliance, where those individuals consuming the most fiber habitually were the least compliant to consuming prebiotic supplements. We found evidence to the contrary and observed an overall missed-dose rate of only 3.6% based on self-report during post-intervention surveys. Importantly, participants were not compensated for their participation in this study, so falsifying compliance surveys was minimally incentivized. We also found that side-effects were not more likely in high baseline fiber consumers, providing further evidence that high fiber consumers were not more likely to skip prebiotic doses. Another concern is that the prebiotic supplements caused behavioral or physiological alterations disproportionately. However, we found that habitual fiber consumption did not decline throughout the study, and no differences in self-assessed Bristol stool hardness were observed. Together, this evidence bolsters our confidence in concluding that the observed negative relationship between baseline fiber consumption and butyrogenic response to prebiotics is real and driven by microbiological factors. We also acknowledge our modest sample size of 28 individuals, and note that some additional relationships may become apparent with greater statistical power. Future and larger prebiotic trials are needed to determine the generality of our observed association between habitual diet and SCFA production.

## Conclusions

Our study provides evidence that the responsiveness of an individual to prebiotic treatment may be predictable from diet and baseline concentrations of SCFA in stool. We emphasize that we do not believe these predictions should be used to modify the diets of individuals who normally consume high levels of fiber or have higher concentrations of SCFA in stool. Ample evidence supports the benefits of both a high fiber diet and adequate microbial SCFA production (71, 98, 99). Rather, we argue our findings support strategies for personalizing microbiota therapy that focus on individual selection. Prior work on prebiotic responses have primarily used human studies with parallel design (30, 32, 39), which do not allow analysis of individualized response across prebiotics. Here we show, we believe for the first time, that the responses within individuals are correlated across at least some pairs of prebiotics, and that average response correlates with host factors. Ongoing prioritization efforts are being used to identify patients most likely to respond to microbiota-directed and microbiota-dependent disease therapies (100–103), and are also being explored to enhance prebiotic therapy (30, 70, 93, 94). Our study here suggests that such efforts to personalize prebiotic supplementation can rely on dietary history and fecal metabolite levels as informative and non-invasive biomarkers of treatment outcome.

## Supporting information

SCFA data

ASV table

Sequencing metadata

Sequencing taxonomy table

## List of Abbreviations

DHQ3: diet history questionnaire, version 3
GCFID: gas chromatograph – flame ionization detector
GLM: generalized linear model
GLMM: generalized linear mixed-effects model
GOS: galactooligosaccharides
SCFA: short-chain fatty acids
USDA: United States Department of Agriculture

## Declarations

### Ethics approval and consent to participate

All recruitment and study procedures involving human subjects were approved by the Duke University Institutional Review Board (Pro00087214). All subjects provided electronic informed consent prior to enrollment.

### Consent for publication

Not applicable.

### Availability of data and materials

The datasets generated and analyzed during the current study, along with analytical code, are either included in the supplement or available in the figshare repository, https://doi.org/10.6084/m9.figshare.c.5405505. The raw sequencing data is available on the National Center for Biotechnology Information (NCBI) Sequence Read Archive (SRA) under the accession PRJNA834995.

## Funding

This study was supported by funding from the Beckman Young Investigator program, the Damon Runyon Cancer Research Foundation, the NASA Translational Research Institute through Cooperative Agreement NNX16AO69A, and NIH 1R01DK116187-01. This work used a high-performance computing facility partially supported by grant 2016-IDG-1013 (HARDAC+: Reproducible HPC for Next-generation Genomics”). ZH was funded by the National Defense Science and Engineering Graduate fellowship. MV was funded by a Postdoctoral Enrichment Program Award from the Burroughs Wellcome Fund. JS and BP were funded by the Duke School of Medicine Medical Scientist Training Program. BP was additionally funded by the Integrative Bioinformatics for Investigating and Engineering Microbiomes (IBIEM) program. No funding body had any role in the design of the study nor the collection, analysis, or interpretation of the data nor in writing the manuscript.

### Authors’ contributions

ZH, MV, JS, LD, and HD conceptualized and designed the study. HD, ZH, SJ, and LD coordinated recruitment and engaged with participants. ZH was blinded to participant identity and study arm assignment where possible. ZH, SJ, ED, and HD processed samples, ran *in vitro* fermentation assays, and analyzed samples via gas-chromatography and sequencing. PHL and BP supported collection and analysis of nutrition data. ZH analyzed all data. ZH and LD wrote the manuscript. All authors read and approved the manuscript.

## Acknowledgements

The authors acknowledge the Stedman Nutrition Center at Duke University for lending kitchen facilities for food-safe repackaging of prebiotics, and Jeffrey Letourneau, Rachael Bloom, and Savita Gupta for assistance with study coordination and logistics.

## Supplementary figures and tables

**Figure S1:**
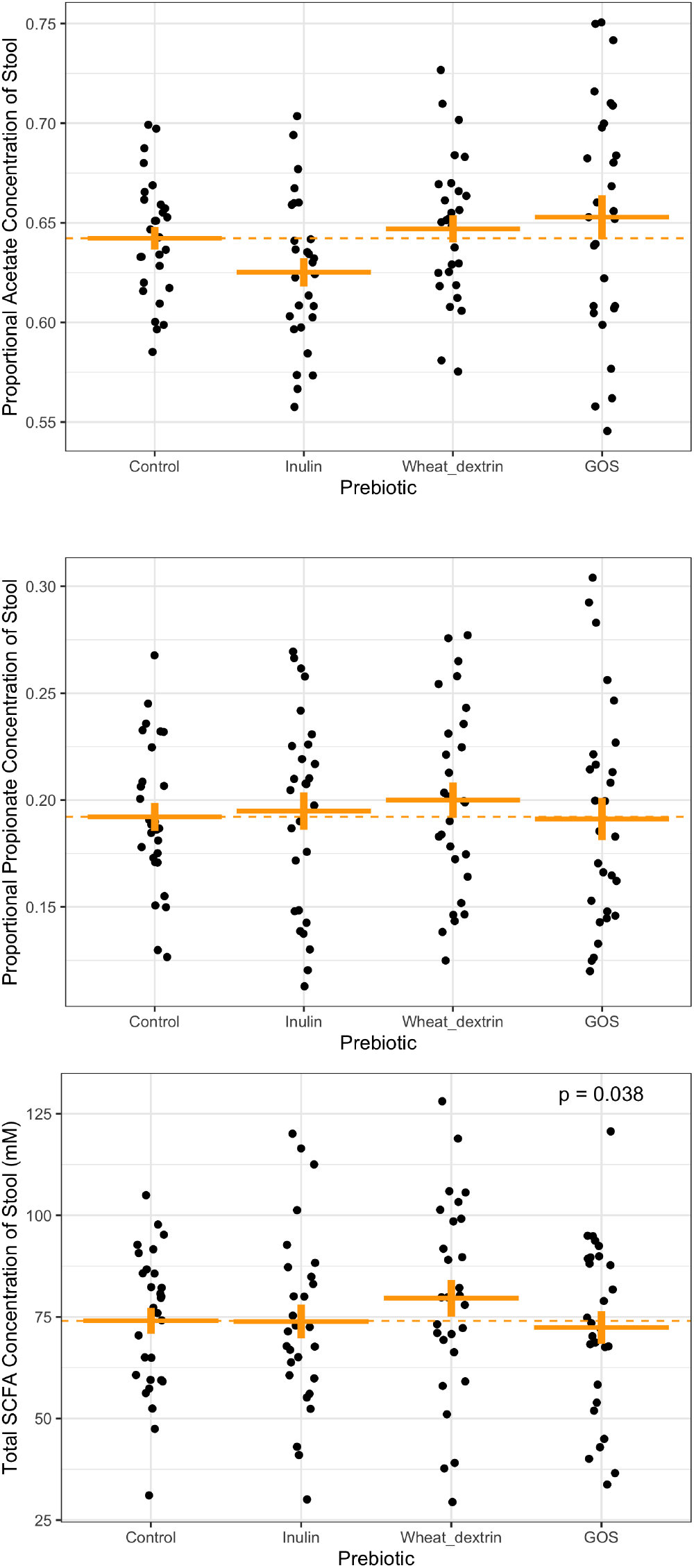
Proportional Acetate (A), proportional propionate (B), and total SCFA (C) concentrations during baseline and prebiotic interventions. All significant differences between prebiotic treatments and control are designated by displayed p values, calculated using a series of generalized linear mixed models. *A proportional SCFA concentration value of 1 = 100%*.

**Figure S2:**
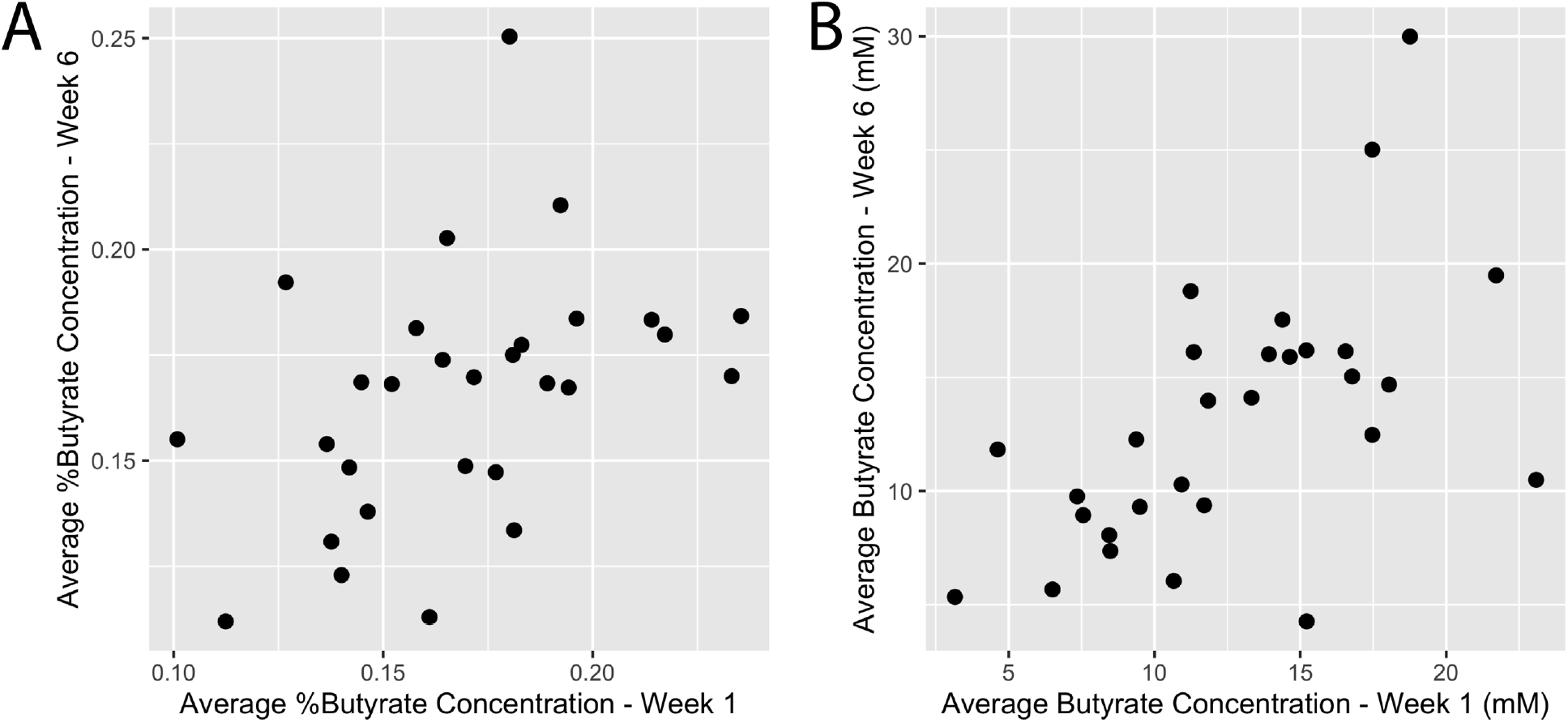
Average proportional butyrate concentration (A) and average butyrate concentration (B) in participant stool during the first and last week of the study. Both proportional concentration and raw concentration were correlated between the first and last weeks (ρ = 0.49, p < 0.007; and ρ = 0.63, p = 0.0003, respectively, Spearman correlations).

**Figure S3:**
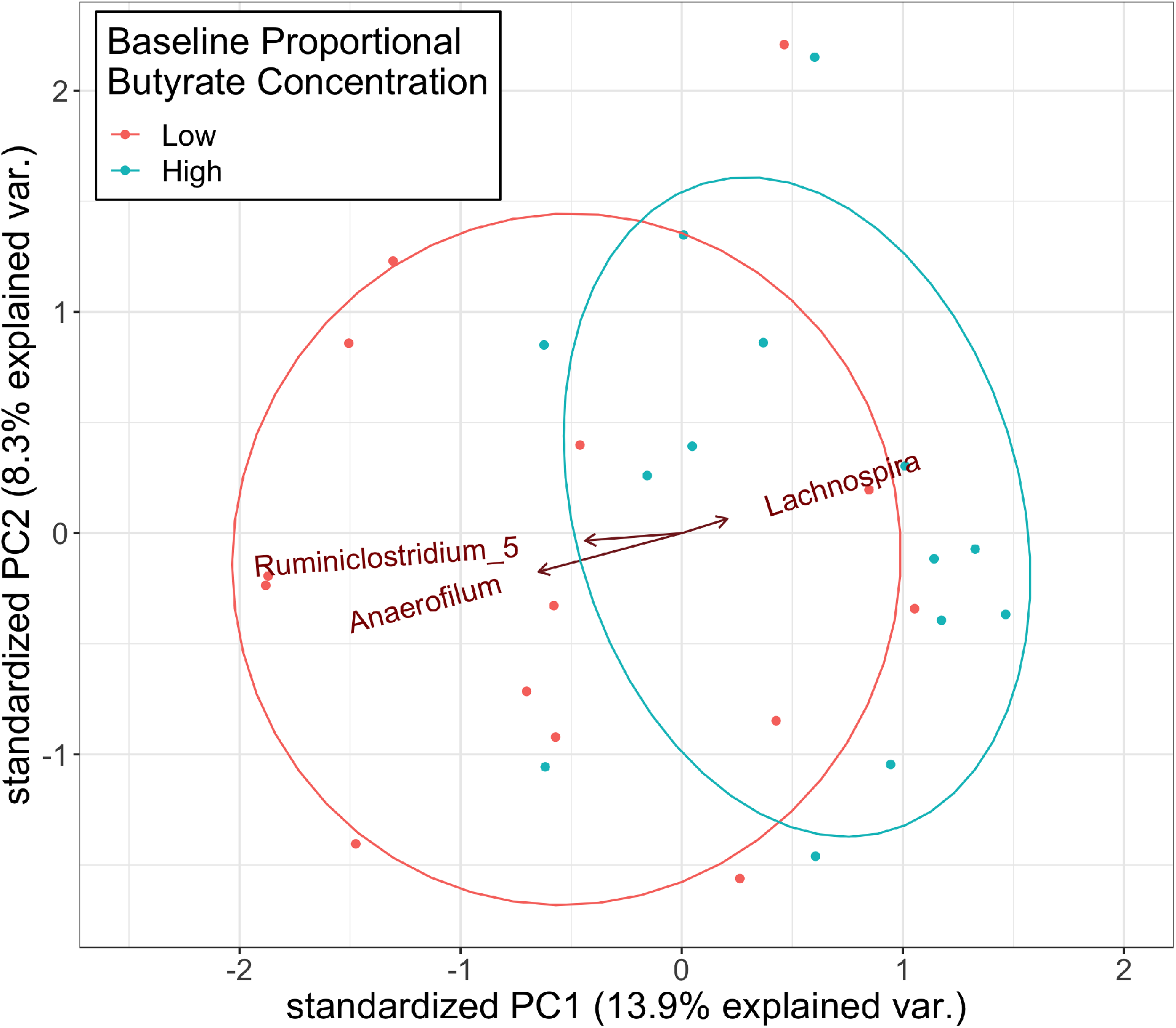
Biplot of stool 16S rRNA gene community composition showing samples (dots) plotted on two principal component axes along with corresponding taxonomic variables (arrows), agglomerated to genus level. Samples are colored by group (high and low; > median baseline %butyrate and <= median baseline %butyrate, respectively) and ellipses represent 95% confidence intervals of the centroid for each group. Centroids are significantly different (p = 0.0022; PERMANOVA using unweighted UniFrac distance). Only significantly differentially abundant genera (Benjamini-Hochberg corrected q < 0.05, ALDEx2) are represented by arrows and labels.

**Figure S4:**
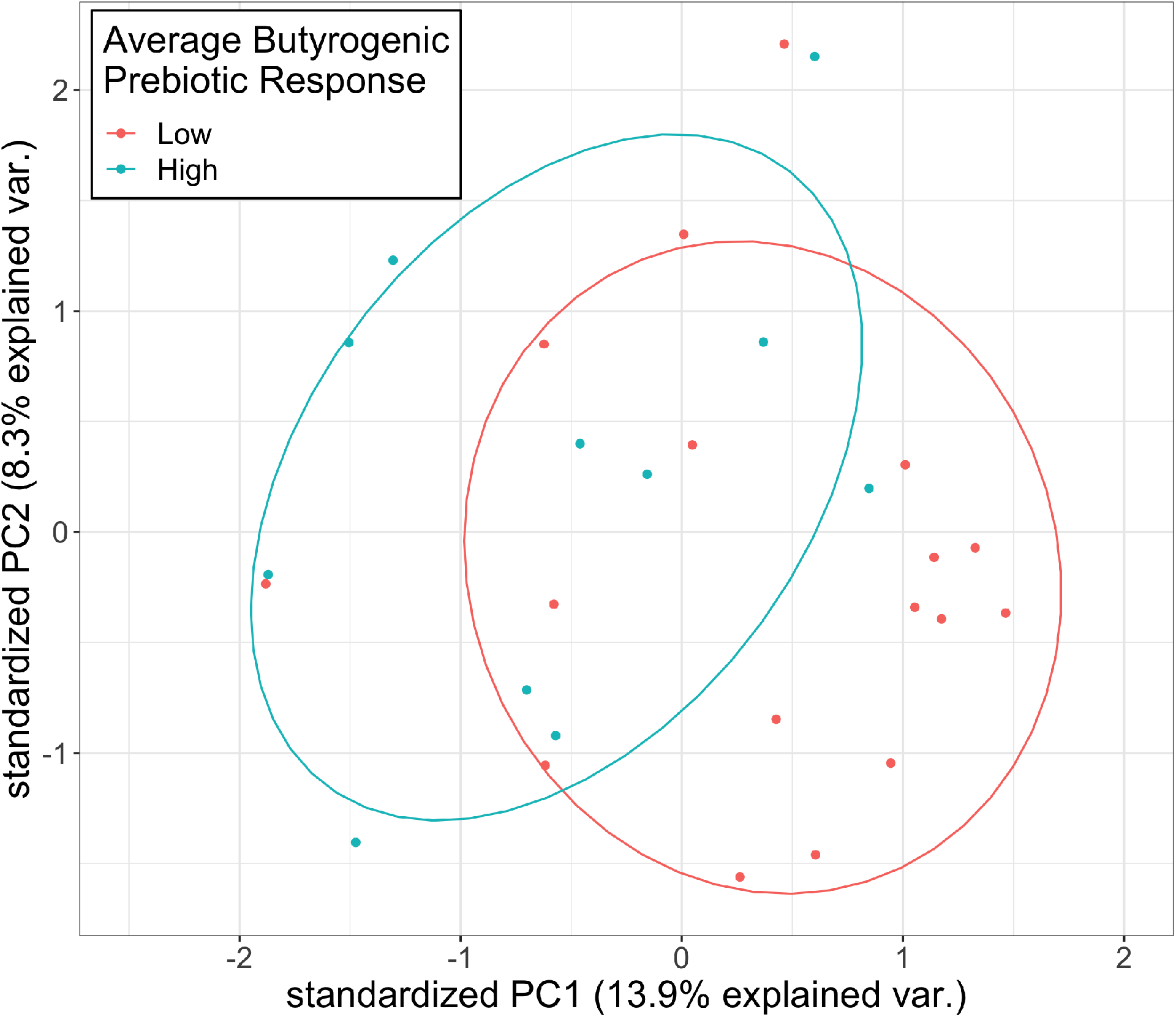
PCA of stool 16S rRNA gene community composition showing samples plotted on two principal component axes. No genera were significantly differentially abundant (Benjamini-Hochberg corrected q < 0.05, ALDEx2), so no variables are plotted. Samples are colored by group (high and low; > 0 average butyrogenic response to prebiotics and < 0 average butyrogenic response to prebiotics, respectively) and ellipses represent 95% confidence intervals of the centroid for each group. Centroids are significantly different (p = 0.028; PERMANOVA using unweighted UniFrac distance).

**Figure S5:**
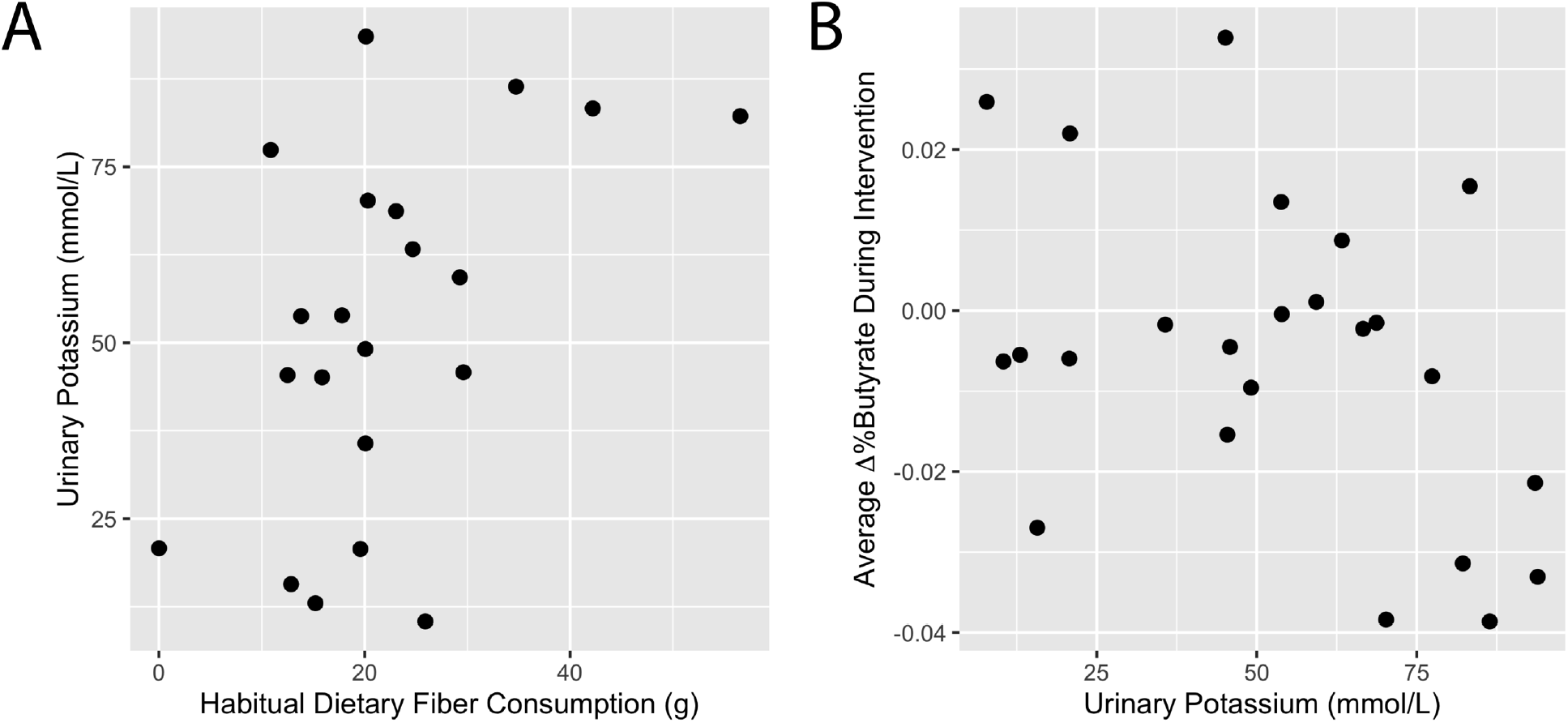
Relationships between baseline urinary potassium and (A) habitual dietary fiber consumption, and (B) average change in proportional butyrate concentration during prebiotic interventions. Baseline urinary potassium was positively correlated with habitual fiber consumption (ρ = 0.47, p = 0.031; Spearman correlation; A) and showed evidence for a negative correlation with average change in proportional butyrate concertation during prebiotic interventions (ρ = −0.38, p = 0.071; Spearman correlation; B).

**Figure S6:**
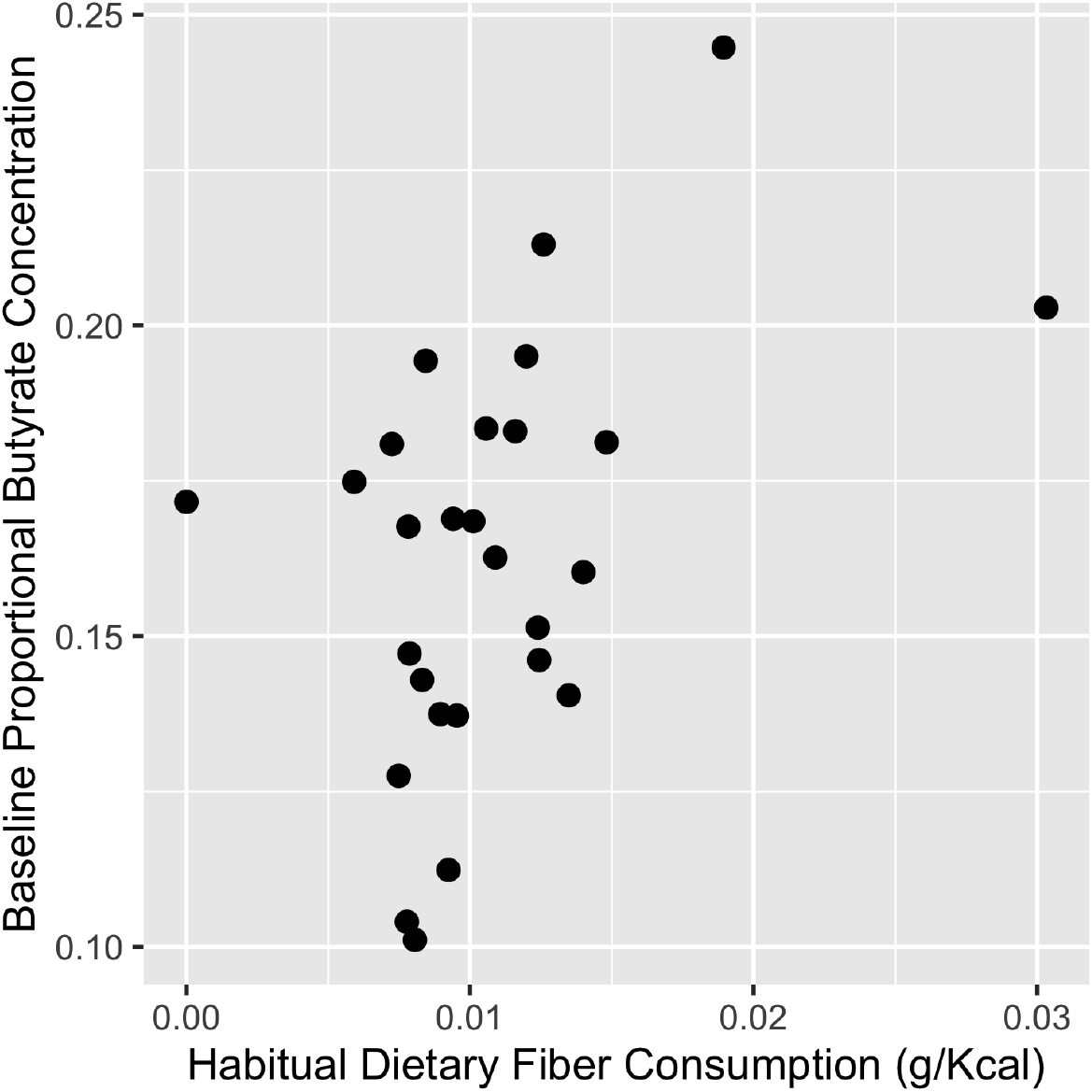
Relationship between habitual dietary fiber consumption and baseline proportional butyrate concentration at week one (prior to study intervention). Fiber consumption was positively associated with baseline proportional butyrate concentration, with 1g fiber/2000 kilocalorie diet associated with an increase in proportional butyrate concentration of 0.010 (p = 0.012, beta regression). A proportional butyrate concentration value of 1 = 100%.

**Figure S7:**
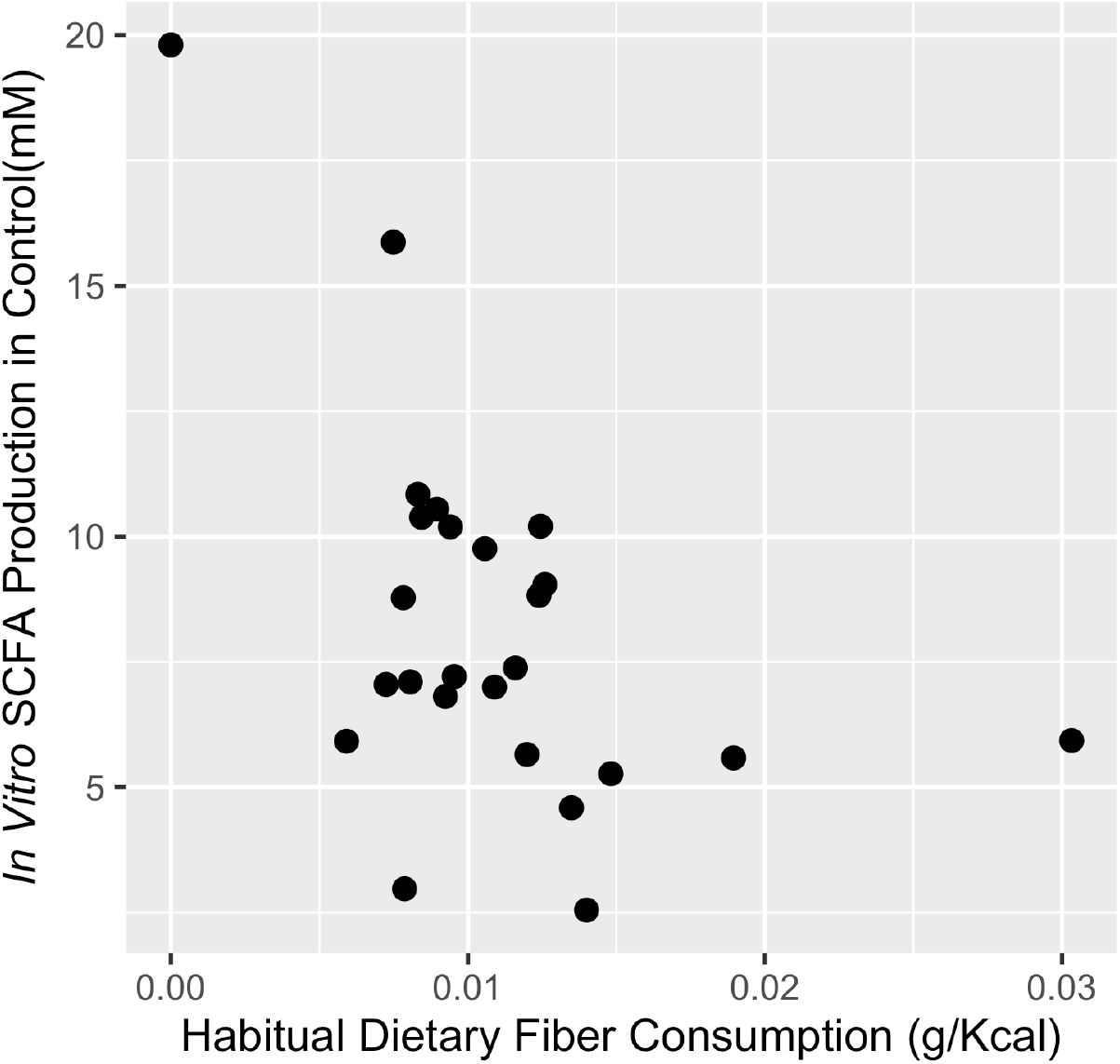
Relationship between *in vitro* SCFA production in prebiotic-free control treatments and habitual dietary fiber consumption. *In vitro* SCFA production was negatively correlated with habitual fiber consumption (ρ = −0.43, p = 0.031; Spearman correlation).

